# Judgement Bias During Pregnancy in Domestic Pigs

**DOI:** 10.1101/2021.08.13.456208

**Authors:** Emily V. Bushby, Sheena C. Cotter, Anna Wilkinson, Mary Friel, Lisa M. Collins

## Abstract

In humans and rats, changes in mood and affect are known to occur during pregnancy, however it is unknown how gestation may influence mood in other non-human mammals. This study assessed changes in pigs’ judgment bias as a measure of affective state throughout gestation. Pigs were trained to complete a spatial judgement bias task with reference to positive and negative locations. We tested gilts before mating, and during early and late pregnancy, by assessing their responses to ambiguous probe locations. Pigs responded increasingly negatively to ambiguous probes as pregnancy progressed and there were consistent inter-individual differences in baseline optimism. This suggests that the pigs’ affective state may be altered during gestation, although as a non-pregnant control group was not tested, an effect of learning cannot be ruled out. These results suggest that judgement bias is altered during pregnancy in domestic pigs, consequently raising novel welfare considerations for captive multiparous species.

## Background

Research investigating the links between pregnancy, affect and cognition is most often carried out with a human-centric focus with studies typically using case studies and cohorts. In humans, changes in affective state during pregnancy are common and alterations in levels of anxiety, depression and cognitive ability have been demonstrated in humans and rodents [1–3]. These changes are often linked to the large and rapid hormone fluctuations that occur during the gestational period [4–5]. Where human subjects cannot be used, rodent models are often employed to experimentally investigate how factors such as diet, enrichment or stress can influence behaviour during pregnancy [6–8]. To infer anxiety and depressive-like behaviours, lab-based behavioural tests, such as a forced swim or open-field test are often used [9]. These studies are conducted under laboratory conditions and are generally aimed at modelling human gestation, rather than investigating how gestation may impact on the rodent itself. Results from both human and rodent studies are varied, however most show that affective state is altered throughout gestation (for review see [2]) and it is clear that pregnancy impacts maternal affective state.

Understanding an animals’ affective state, or emotion and mood, is a key component of animal welfare [10]. Affective state can influence and alter cognitive processes, such as judgement, [11–12] which may then be used to infer and understand an animals’ affective state. Cognitive bias or judgement bias is the influence of affect on information processing, with more content individuals likely to make positive assumptions about ambiguous stimuli [13]. Judgement bias tests have been used to assess changes in affective state in a range of species, including pigs, dogs, honeybees and European starlings [14–17]. Research typically focuses on the impact of external stimuli on judgement bias; this is likely to act via alteration to the internal, physiological environment ultimately resulting in changes in behaviour and judgement bias [11; 18–19]. As such, we would expect internal stimuli, such as physiological changes, would also impact judgement bias directly even in the absence of external influences. Pregnancy is one of the biggest physiological changes a mammal may experience, involving major hormonal and cognitive adjustments [20–21], yet little is known of how information processing and affective state may change in relation to pregnancy in animals.

The domestic pig (*Sus scrofa domesticus*) has been used as a human model in a wide range of medical research such as infectious disease [22], nutritional [23] and neurological studies [24]. Pigs allow for longer lifespan studies and are more similar to humans than other laboratory species, such as rodents [25–26]. More commonly, pigs are farmed around the globe for meat production. Modern intensive farming systems have been designed to produce food as quickly and cost efficiently as possible, and research is continually ongoing to understand how animal welfare can be optimised within these systems. Despite many studies on the behavioural and welfare needs of sows during pregnancy [27–30], only one study used a specific judgement bias task to assess affective state in gestating sows. This study focused on whether judgement bias could be used as a welfare indicator in gestating sows, finding that group-housed, gestating sows can learn a go/no-go judgement bias task and individual affective states can differ despite experiencing the same management conditions [31]. However, this study did not investigated how gestation itself influenced judgement bias. More recently another study showed that gestating gilts that were classified as ‘friendly’ visited an electronic sow feeder more often than individuals that were classified as ‘fearful’ [32]. The authors hypothesised that this feeding behaviour may be similar to a judgement bias task and that the friendly individuals may have been more optimistic. However, again this study did not investigate how gestation itself influenced judgement bias.

We investigated how gestation may alter judgement, and therefore affective state, in domestic pigs. We compared within- and between-individual affective state, as measured by a spatial judgement bias test, before mating, and during early and late pregnancy. We hypothesised that within-individual judgement bias would be more pessimistic during pregnancy than prior to mating, leading to an increase in latency to approach ambiguous cues throughout pregnancy. This is the first study to our knowledge to investigate the possible impact of gestation on judgement bias in domestic pigs.

## Methods and materials

This work was carried out between July and October 2015 (replicate one) and between January and July 2017 (replicate two) on a pig farm in the UK.

### Animal housing and husbandry

20 gilts (primiparous female pigs; N=10 for each replicate) were selected based on age and time until first mating. Using gilts allowed for training time before gestation, as there is limited time between pregnancies once a sow has begun breeding. The average age of all 20 pigs on day one of training was 241.7 (SE: 3.56) days. Replicate two contained one Duroc and three Landrace pigs, the breed of all other individuals was Large White. Pigs were housed in pens of five or six animals, each pen (4.67m x 5.35m) contained a sheltered sleeping area with straw bedding (2.70 x 4.67m) and a run partially exposed to outdoor elements, such as wind and natural light (2.65 x 4.67m). A standard lactating sow ration was fed once a day before mating and throughout gestation; there was continuous access to water and natural lighting. During the course of the study the animals remained within the same groups and pens to keep the external environment as controlled as possible throughout. The study pigs were able to interact with pigs in the pen next door via the gate and animals in the neighbouring pens may have been moved/changed. Due to involvement in a separate study, replicate one pigs received Regumate® (containing a steroidal progestin) orally with feed 23 days before planned estrus to allow for synchronised farrowing. As of June 2020, no previous research was found investigating possible effects of Regumate® on affective state or behaviour of pigs. Due to this research taking place on a working farm, it was not possible to test a non-pregnant control group

### Judgement bias

All pigs were habituated to the test arena in groups for two to three sessions, and then individually to habituate the pigs to eating from the bowl which was placed in the centre of the test arena. Following this, individuals were trained to associate the bowl location with a positive (P) and a negative (N) outcome. When in the P location, the bowl contained a small amount of chocolate raisins (replicate 1) or sugar-coated chocolates (replicate 2) and when it was in the N location, the bowl contained unpalatable food (bitter tasting coffee beans) to discourage the pigs from approaching this location. The pigs were trained to discriminate between these reference locations by alternating P and N trials. Latency to reach the bowl was recorded using video cameras and was then used as a metric to assess whether each individual had learned the discrimination. Each trial was 30 seconds in duration. Correct responses were recorded when the subject approached and touched their nose to the bowl during the positive (P) trials; during negative (N) trials, a correct response was recorded when the individual did not approach the bowl within 30 seconds. The location of P and N was counterbalanced across individuals. For both replicates a criterion of 70% correct responses in the last 20 trials was required before moving onto the testing phase. Per individual, forty-four training trials were conducted during replicate one and sixty-two for replicate two. Replicate two required more training trials due to the pigs being slower to differentiate between the positive and negative locations. Five pigs from replicate one failed to meet this criterion and were removed from the study. Two pigs from replicate 2 did not meet this criterion. The analysis represents only those 13 that met the learning criterion.

Each testing session comprised two sets of nine trials carried out on the same day, involving five different bowl locations; the P and N reference locations and three intermediate ambiguous probes: near positive (NP), middle (M) and near negative (NN). Only one bowl was in the arena during each trial. The ambiguous probes placed in predetermined equidistant positions (0.74m) and were not reinforced (i.e., they were left empty). They were presented in a pseudo-randomised order and interspersed among training trials. All ‘during pregnancy’ testing sessions were preceded by five ‘reminder’ training trials the day before testing. Each pig was tested three times: before gestation (1-2 weeks before mating); early gestation (4 weeks after mating); and late gestation (10-11 weeks after mating). One pig in replicate two was not tested before gestation and was only tested in the early and late test phases.

### Statistical analysis

All data were analysed in R version 3.4.1 using general linear mixed effects models with the *lmer* function in the package *lme4* [33]. To test the effects of gestation time on cognitive bias, the response variable was *time taken to approach* the presented probes; fixed explanatory effects were *probe location,* coded as a continuous variable from positive (1) to negative (5) with ambiguous locations at points 2, 3 and 4; and *gestation time* coded as a factor with three levels (pre, early and late gestation). *Probe location squared* was included as initial data exploration suggested curvature in the fits. Interactions between *gestation time* and *probe location* and *probe location squared* were also included.

To find the most appropriate structure for the random model, we compared eight models: two intercept only models and six combinations of random intercept and slope models such that random intercepts were fitted for each pig at each experimental timepoint (or for each pig independent of experimental replicate), with variation allowed between gestation times and the shape of the curve was allowed to vary between pigs (Table 1).

**Table 1:** Statistical model details. Random models with fixed slopes (models 1 and 2) or slopes allowed to vary across probe location (models 3-8), with experimental replicate included (models 2,4,6 and 8) or not (models 1,3,5 and 7).

| Model | Random slope | Random intercept |
| --- | --- | --- |
| 1 | 1 | Gestation time:Pig ID |
| 2 | 1 | Replicate/Gestation time:Pig ID |
| 3 | Location | Gestation time:Pig ID |
| 4 | Location | Replicate/Gestation time:Pig ID |
| 5 | Location<sup>2</sup> | Gestation time:Pig ID |
| 6 | Location<sup>2</sup> | Replicate/Gestation time:Pig ID |
| 7 | Location+Location<sup>2</sup> | Gestation time:Pig ID |
| 8 | Location+Location<sup>2</sup> | Replicate/Gestation time:Pig ID |

The Akaike Information Criteria (AIC) values for all models were compared using the *model.sel* function in the *MuMIn* package [34]. In each case the residuals of the final minimal model were visually assessed for deviations from normality. For the final models, predicted fits were produced using the *predict* function in base R. R^2^ values for each model were calculated using the *r.squaredGLMM* function in the *MuMIn* package [34]. For every model, the general pattern of results was robust, with the different random models only affecting the predictions very slightly. The best model is reported in the main text, and the results and corresponding figures for the two models where AIC comparison had delta < 2 are reported as supplementary information.

## Results

### Judgement Bias

The pigs’ responses to ambiguous locations in the cognitive bias test changed throughout gestation (Tables 2, 3; Figure 1). Pigs consistently approached the positive probe quickly and the negative probe slowly (or not at all), getting generally slower during gestation (Figure 1). However, whilst the mean speed of approach was fairly linear between positive and negative pre- and early gestation (Figure 1a,b), by late gestation, pigs showed a shift towards pessimism, such that the positive probe continued to be approached quickly but ambiguous probes were approached more slowly (Figure 1c).

**Figure 1:**
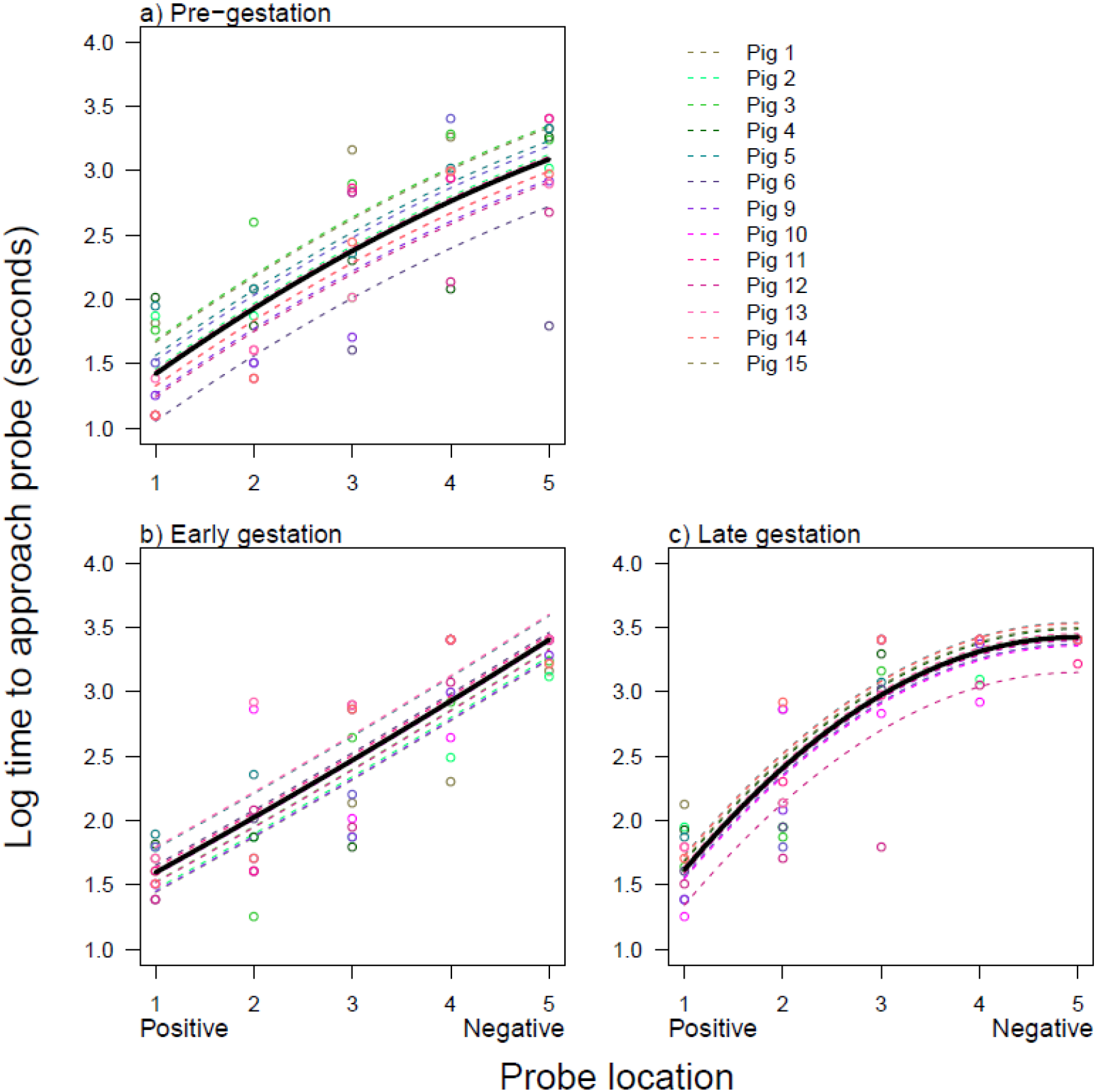
The time to approach each location at three stages of gestation. Log time taken to approach each location for pigs at three different stages of the pig’s 16-week gestational period; a) pre-gestation, b) early gestation (5 weeks) and c) late gestation (10-11 weeks). The open circles are raw datapoints and the lines are model predictions from the minimal adequate model fixed to the level of experimental replicate 1. Results from model 2 are shown, where the intercept is allowed to vary for each pig at each gestation time, within each replicate. Pigs 1-5 are from replicate 1 and pigs 6-15 are from replicate 2.

**Table 2:** Table of candidate LMERs. Table of candidate LMERs explaining time to approach the probe in relation to the interaction between the location of the presented probe and the gestation time for pigs that reached the 70% learning criterion only (n=13). Each model retained all fixed terms (Location*Gestation time+Location^2^*Gestation time) with only the random model varying. *Model* corresponds to the random model listed in Table 1, *AIC_c_* = corrected Akaike Information Criteria values; Δ *AIC_c_* = difference in *AIC_c_* values between the best model (lowest *AIC_c_*) and the given model; *w* = Akaike weights; *r*^2^ (F only) = *r*^2^ for the fixed model only, *r*^2^ (F+R) *r*^2^ for the fixed plus random model.

| Model | df | $AIC_c$ | $\Delta AIC_c$ | w | $r^2$ (F only) | $r^2$ (F+R) |
| --- | --- | --- | --- | --- | --- | --- |
| 1 | 12 | 215.2 | 0.00 | 0.580 | 0.751 | 0.805 |
| 2 | 11 | 216.9 | 1.72 | 0.245 | 0.748 | 0.806 |
| 3 | 16 | 219.7 | 4.50 | 0.061 | 0.751 | 0.805 |
| 5 | 13 | 219.8 | 4.57 | 0.059 | 0.751 | 0.805 |
| 7 | 13 | 221.2 | 6.01 | 0.029 | 0.750 | 0.818 |
| 4 | 22 | 222.2 | 7.02 | 0.017 | 0.738 | 0.810 |
| **6** | 16 | 224.1 | 8.89 | 0.007 | 0.740 | 0.817 |
| **8** | 16 | 227.5 | 12.35 | 0.001 | 0.729 | 0.833 |

**Table 3:** Results of the best supported statistical models. Minimum adequate linear mixed effects model for the effects of probe location and gestation time on the time taken for pigs to approach the probe under testing, for pigs that reached the 70% learning criteria only (n=13). The results equate to the best supported random models.

|  | Model 1 |  |  | Model 2 |  |  |
| --- | --- | --- | --- | --- | --- | --- |
| Term | DF | F | P | DF | F | P |
| **Location** | 1, 141 | 62.96 | <0.001 | 1, 141 | 62.96 | <0.001 |
| **Gestation time** | 2, 168 | 2.03 | 0.134 | 2, 167 | 2.04 | 0.133 |
| **Location<sup>2</sup>** | 1, 141 | 9.57 | 0.002 | 1, 141 | 9.57 | 0.002 |
| **Location: Gestation time** | 2, 141 | 6.07 | 0.003 | 2, 141 | 6.07 | 0.003 |
| **Location<sup>2</sup>: Gestation time** | 2, 141 | 6.16 | 0.003 | 2, 141 | 6.16 | 0.003 |

All models retained all interactions and gave qualitatively similar results. The best model was model 1, where the intercept was allowed to vary for each pig at each gestation time (Table 1). However, the result for model 2, where the intercept was allowed to vary for each pig at each gestation time, within each replicate, was equally well supported (delta AIC <2; Table 2, Figure S1).

## Discussion

Commercially farmed breeding pigs often experience multiple consecutive pregnancies throughout their lifespan. In livestock species judgement bias tasks have most commonly been used to assess the impact of external factors on affective state. However, internal factors, such as the large physiological changes as associated with pregnancy, also have the potential to influence affective state and therefore judgement bias. The aim of this study was to assess judgement bias in domestic pigs throughout gestation. It was hypothesised that the gilts would be more pessimistic during pregnancy than prior to mating, as indicated by an increase in latency to approach the ambiguous cues. Our results showed this to be the case, with the gilts taking longer to approach the ambiguous locations in the later stage of gestation than before mating which indicates that judgement bias changed as gestation progressed. This was most apparent at the middle and most ambiguous location (Figure 1) and suggests the pigs were more pessimistic during the late gestational stage. Crucially, the latency to reach the positive location did not vary markedly throughout gestation, showing that other changes, for example, increase in weight, did not affect response latencies (Figure 1). Thus, these results show increased pessimism during the late stage of pregnancy, despite the fact that the immediate external environment remained constant. This infers that, alongside external factors, internally-driven factors can also influence judgement bias and affective state in domestic pigs.

The possibility that pigs’ judgement bias may change from a positive to a more negative state during the late stage of gestation suggests that the pigs’ welfare needs may change too. This highlights the importance of considering the impact of large physiological changes, such as pregnancy, on animal welfare. This study may have implications, not only for the welfare of farmed animals that experience gestation, but also for research into affective state during pregnancy in other captive multiparous mammalian species, including how this may impact cumulatively across the life course on their health and welfare. For example, in humans, multiparous women appear to be more at risk and have a different pattern of anxious or depressive symptoms compared to primiparous women [35–36]. In humans, hormone fluctuations and other physiological changes throughout pregnancy are often correlated with changes in mood and affective state [4–5]. Pigs are frequently used as models for humans in medical and pharmaceutical studies [22–23; 39–40], so it is possible that a change in affective state during gestation may be caused by comparative mechanisms, however, further research is required to validate this.

Alongside this interesting result, there are some limitations to take into consideration. Previous studies have shown that multiple testing time points can result in an increase in pessimistic responses [37–38] and this increase in latencies during the later testing phases is similar to what was found in this study. As it was not possible to test a non-pregnant control group, this effect of learning cannot be ruled out. However, the effects of gestation represent a plausible driver for the changes in affect we report as previous research in rodents and humans has shown that mood and affective state can vary throughout gestation [1–3]. Future studies should consider the role of learning by including a non-pregnant control group. There were also some differences between replicates, such as one replicate receiving Regumate®, and different rewards being used. Despite this, the effect of replicate on the data was marginal (Figure 1), showing that the change in judgement bias over the course of pregnancy was robust and not influenced by these differences between replicates.

In conclusion, this study shows that judgement bias in farmed domestic pigs may change with stage of gestation, inferring that internally driven stimuli can directly affect judgement bias without external influence. This study raises novel welfare considerations for captive multiparous species and provides a basis for future research into the effect of gestation on judgement bias in non-human animals.

## Acknowledgements

We would like to thank the farm staff for their help and cooperation.

## Data accessibility

Data is available as supplementary material on Dryad (Doi to be confirmed)

## Competing interests

The authors declare no competing interests

## Author contributions

EVB carried out data collection. LMC conceived of the study; SC carried out statistical analysis; EVB, SC, AW, MF and LMC assisted with study design and coordination, drafting the final manuscript and gave final approval for publication.

## Ethical statement

The University of Lincoln, College of Science Ethics Committee approved this study. COSREC189, COSREC262.

## Funding

EVB, SC and AW had no funding. LMC and MF were funded by the Biotechnology and Biological Sciences Research Council grant BB/K002554/2

## Supplementary Material

**Figure S1:**
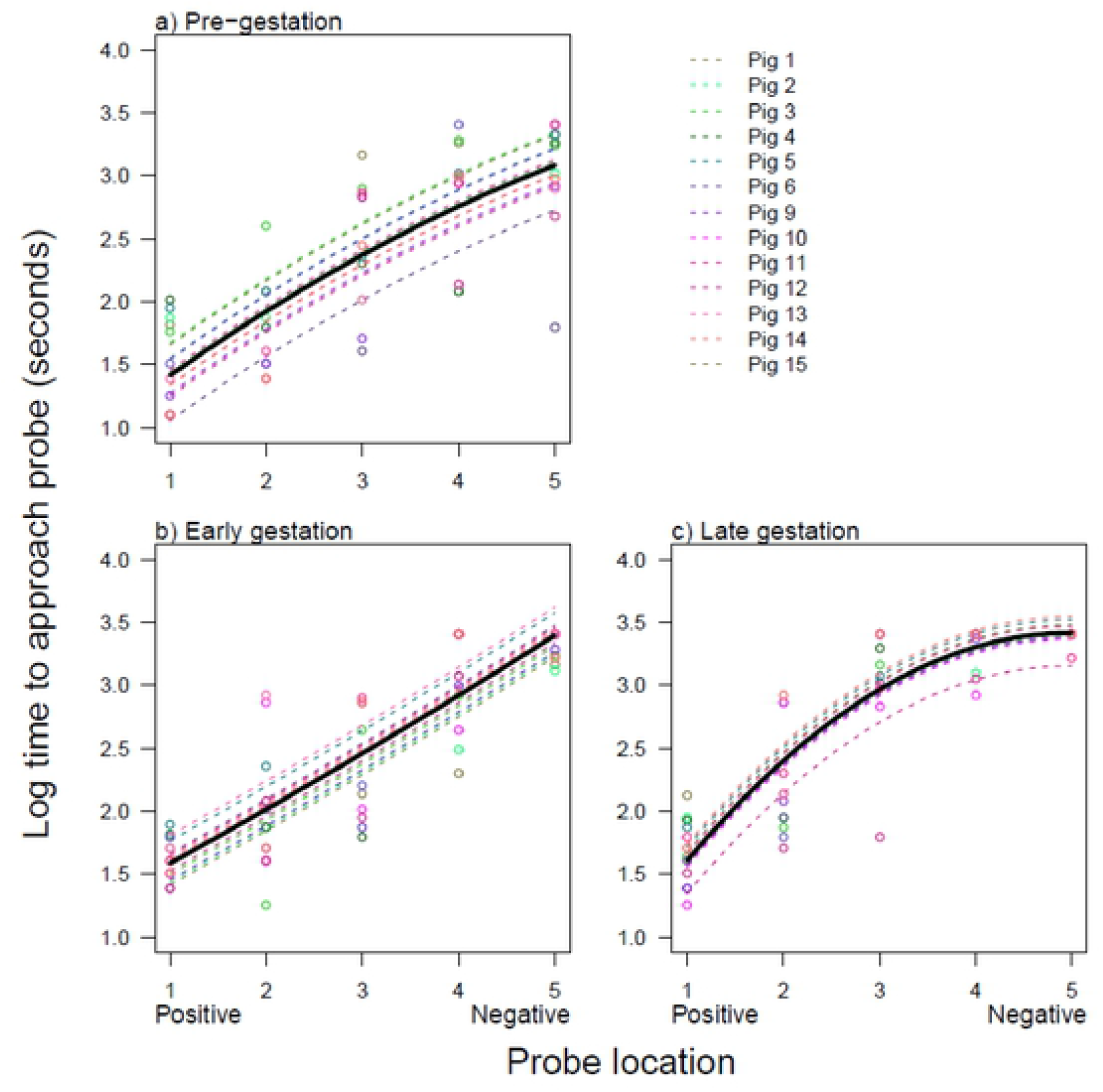
The latencies for all 13 pigs to approach each location during three stages of gestation. Log time taken to approach each location for pigs at three different stages of the pig’s 16-week gestational period; a) pre-gestation, b) early gestation (5 weeks) and c) late gestation (11 weeks). The open circles arc raw data points and the lines arc model predictions from the minimal adequate model fixed to the level of experimental replicate 1. Results from model 1 arc shown, where the intercept is allowed to vary for each pig at each gestation time. Only the 13 pigs that had >70% correct responses during the learning phase arc included.

## Notes

### Competing Interest Statement

The authors have declared no competing interest.

